# U1 AMO (antisense morpholino oligo) disrupts U1 snRNP structure to promote intronic premature cleavage and polyadenylation (PCPA)

**DOI:** 10.1101/2023.02.24.529985

**Authors:** Qiumin Feng, Zejin Lin, Yanhui Deng, Yi Ran, Andy Peng Xiang, Congting Ye, Chengguo Yao

## Abstract

Functional depletion of U1 snRNP with a 25 nt U1 AMO (antisense morpholino oligonucleotides) may lead to intronic premature cleavage and polyadenylation (PCPA) of thousands of genes, a phenomenon known as U1 snRNP telescripting; however, the underlying mechanism remains elusive. In this study, we demonstrated that U1 AMO could disrupt U1 snRNP structure both in vitro and in vivo, thereby affecting U1 snRNP/RNAP polymerase II (RNAPII) interaction. We further showed that U1 AMO treatment might promote RNAPII disassociation with pre-mRNA in an RNA pull-down assay. By performing ChIP-seq for phosphorylation of Ser2 (Ser2P) and Ser5 (Ser5P) of the C-terminal domain (CTD) of RNA polymerase II (RNAPII), we showed that transcription elongation was disturbed upon U1 AMO treatment, with a particular high Ser2P signal at intronic cryptic polyadenylation sites (PASs). In addition, we showed that core 3’ processing factors CPSF/CstF are involved in the processing of intronic cryptic PAS. Their recruitment accumulated toward cryptic PASs upon U1 AMO treatment, as indicated by ChIP-seq and iCLIP-seq analysis. Furthermore, we showed that most of these PCPAed transcripts could be exported to cytoplasm and have the potential to be translated. Conclusively, our data provide more insight into U1 snRNP telescripting, and suggest a common theme that modulation of transcription elongation may be an important mode for the regulation of mRNA polyadenylation.

## Introduction

Full-length transcription of thousands of genes requires U1 snRNP to inhibit mRNA 3’processing at cryptic intronic polyadenylation sites (PAS) of RNA polymerase II (RNAPII) transcripts, which has been termed as ‘U1 snRNP telescripting’ (1–4). Several possible molecular mechanisms have been proposed since its discovery. First, the ‘U1-CPAF’ model states that U1 snRNP transiently associates with CPAF (cleavage and polyadenylation-associated factors), and the 25 nt U1 AMO (antisense morpholino oligonucleotides) targets the 5’ end of U1 snRNA, leading to activation of CPAF near cryptic PAS while maintaining the integrity of U1 snRNP structure (4–7). This activation is partially achieved by remodeling the CPAF components. Notably, most conclusions are based only on quantitative mass spectrometry analysis. Second, the ‘U1-FUS co-inhibition’ model suggests that FUS protein may also play a role in U1 snRNP telescripting, and FUS and U1 snRNP most likely function as a complex (8–10). Although FUS potentially provides another new mechanism for U1 snRNP telescripting, it has only been examined in specific neuronal cell lines and whether FUS plays a general role in U1 snRNP telescripting remains unknown. Third, we previously hypothesized a ‘U1 snRNP steric hindrance’ model (11–13), which suggests that U1 snRNP may inhibit the assembly of a fully functional 3’ processing complex on cryptic PAS. An obvious limitation of this study is that the results from specific RNA substrate might not reflect the general mechanisms. It must be pointed out that co-transcriptional mRNA 3’ processing has yet to be accounted for in any of the above models (13). Additionally, although previous RNAPII ChIP-seq analyses have suggested that U1 snRNP telescripting is correlated with transcription (3,4), the underlying mechanisms remain elusive.

To extend our U1 snRNP steric hindrance model, we investigated the effect of U1 AMO on U1 snRNP, and unexpectedly found that it could disrupt U1 snRNP structure both in vitro and in vivo. We further provided evidence that U1 AMO may directly impact the association of U1 snRNP and RNAPII, thereby impacting the transcription activity of the latter. This U1 AMO-induced premature transcription termination (PTT) might cause premature cleavage and polyadenylation (PCPA). Overall, our data provide an up-to-date understanding of U1 snRNP telescripting.

## Materials and Methods

### Cell culture and oligonucletide/plasmid transfection

HeLa cells were cultured in DMEM supplemented with 10% FBS. Antisense morpholino oligonucleotide (AMO) transfections were performed as previously described using a Bio-Rad Gene Pulser (2,11,12). The working AMO concentration was 10 μM. After transfection, cells were cultured for 12 h or other indicated time points (0h, 3h,6h,9h and12h in Figure 3D). The sequence of the 25-mer U1 AMO is 5’ -GGTATCTCCCCTGCCAGGTAAGTAT-3’. The sequence of control AMO is 5’- CCTCTTACCTCAGTTACAATTTATA-3’. AMOs were ordered from Gene Tools. Transfection of luciferase reporter plasmid pPASPORT constructs was carried out using Lipofectamine 2000 (Thermofisher) according to the user manual. Transfection of siRNAs was carried out using Lipofectamine RNAimax (Thermofisher). The siRNA target sequences were listed in Supplemental Table 7.

### MS2-tagged RNA affinity purification of mRNA 3’ processing complex

To purify the mRNA 3’ processing complex active in the processing of intronic cryptic PAS, we employed a well-established MS2-tagged RNA affinity purification method (14,15). Briefly, three copies of MS2 sequences were inserted upstream of the *nr3c1* intronic PAS (wild type and mutant sequences were listed in Supplemental Table 7) and RNA was synthesized by in vitro transcription (T7 RNA polymerase). The MS2-MBP fusion protein was prepared from E. coli strain BL21 (DE3) and subsequently purified with amylose beads (NEB). The RNA affinity purification of mRNA 3’ processing complex was carried out essentially as described before (15). Protein pellet were analyzed by mass spectrometry to identify 3’ processing factors, or by SDS-PAGE and silver staining or Western blotting analyses. Similar MS2-tagged RNA affinity coupled with Western blotting analysis was performed for Figure 3A. The working AMO concentration for in vitro assay was 10 μM.

### Immunoprecipitation (IP)

Antibodies were coupled to protein A/G Dynabeads (Bimake). A 500 μl mixture containing 150 μl of HeLa nuclear extracts (NE), 150 μl of dilution buffer (20 mM Hepes, pH 7.9; 100 mM KCl; 500 μM ATP; 3.2 mM MgCl2, and 20 mM creatine phosphate) was incubated for 10 min and 30°C. After incubation, mixtures were spun at 4°C for 5 min at 13,000×g. Supernatants were added to 250 μl of IP buffer (1X PBS, 0.1% Triton X-100, 0.2 mM PMSF, protease inhibitor EDTA-free), spun at 4°C for 5 min, and added to 40 μl antibody-coupled beads. After rotation overnight at 4 °C, six washes (0.8 mL each) were performed by using wash buffer (1X PBS containing 0.1% Triton X-100, 0.2 mM PMSF, pH 8.0). Proteins were eluted by SDS sample loading buffer, followed by Western blotting analysis. For RNA IPs, RNAs were extracted by Trizol reagents (Thermofisher), Northern blot analysis of U1 snRNA was performed using 5’-radiolabeled DNA probes (Sequence is listed in Supplemental Table 7) in ultrasensitive hybridization Buffer (Thermofisher). The radioactivity signals were analyzed by PhosphorImager (Typhoon FLA 7000).

### Gel shift assays

Recombinant GST-U1-70K, GST-U1A, and GST-U1C proteins were prepared from E. coli strain BL21 (DE3) and purified with ProteinIso ^®^ GST Resin (Transgen). Gel shift assays were performed following a reported protocol (16). Briefly, P^32^ labeled RNAs (detectable amount, about 0.2 μM) with the indicated amount of recombinant fusion protein and AMO/DNA in 15 μl of binding buffer [10 mM Hepes (pH 7.9), 50 mM NaCl, 0.5 mM MgCl2, 0.1 mM EDTA, 5% glycerol, 1 mM ATP, 10 mM creatine phosphate, 5 mM β-mercaptoethanol, 0.25 mM PMSF, 0.7 μg of Escherichia coli tRNA, and 1.4 μg of BSA] were incubated at 25°C for 10 min. Samples were further loaded onto 5% non-denaturing PAGE gels. Gel electrophoresis was carried out at 10 mA for 1 h in 1XTBE. The radioactivity signals were analyzed by PhosphorImager (Typhoon FLA 7000).

### DNA-biotin based pull-down assay

Biotinylated DNA oligos were ordered from SYNBIO technologies (Sequences were listed in Supplemental Table 7). Biotinylated DNAs were first bound to the streptavidin beads, and then were incubated with HeLa NE in the polyadenylation conditions for 10 min. After biotin-streptavidin binding and washing, pull-down samples were heated at 95°C for 5 min in 1XSDS PAGE sample loading buffer. The eluted samples were further subject to Western blotting analysis.

### In vitro mRNA 3’ processing assays

In vitro cleavage/polyadenylation and in vitro cleavage assays were performed using P^32^-radiolabeled RNA sequences and HeLa NEs following the standard protocol as described elsewhere (12,14,17–19). Briefly, polyadenylation reactions contain 8 pmol radiolabled RNA/ml reaction, 40% NE, 8.8 mM HEPES (pH 7.9), 44 mM KCl, 0.4 mM DTT, 0.7 mM MgCl2, 1 mM ATP, and 20 mM creatine phosphate. In cleavage reactions, ATP was excluded and 0.2 mM Cordycepin (Sigma), 2.5% PVA, and 40 mM creatine phosphate were added. HeLa NEs were purchased from IPRACELL.

### Luciferase reporter assay

HeLa cells were harvested after 24 h of transfection with pPASPORT plasmids. Luciferase activity was measured using Promega Dual-Luciferase Reporter kit and Berthold Sirius detection system.

### Western blotting

The primary antibodies for 3’ processing factors were purchased from Bethyl Laboratories. Other primary antibodies were purchased from Abcam or Santa Cruz Biotechnology (Cat. Number available on request). For secondary antibodies, we used HRP-conjugated anti-mouse/rabbit (Sigma). ECL western blotting system (Thermofisher) was used to detect the signals.

### Nuclear/Cytoplasm Extraction

A Nuclear/Cytosol Extraction Kit (Abcam) was used to prepare nuclear/cytoplasm proteins/RNAs for Western blotting analysis or 3’-seq.

### Immunostaining

Immunostaining was performed using standard procedures (https://www.abcam.cn/protocols/immunocytochemistry-immunofluorescence-protocol). Briefly, control or U1 AMO transfected HeLa cells were grown on a sterile glass coverslip and fixed using paraformaldehyde. After washing and cell permeabilization, primary antibodies (U1C, sc-101549, Santa Cruz) were incubated with cells overnight at 4°C. Secondary antibodies were ordered from Beyotime Biotechnology. The fluorescence signals were detected with a Confocal Laser Scanning Microscopy (LSM880).

### 3’-seq, mRNA-seq, Ribo-seq, ChlP-seq, and iCLIP-seq

3’-seq was carried out using QuantSeq Rev 3’ mRNA sequencing library prep kit (Lexogen). Downstream sequence analysis was carried out using our recently published pipeline QuantifyPoly(A) (20).

Preparation of mRNA-seq/Ribo-seq libraries, sequencing, and analysis of sequence data was performed according to Illumina and Novogene’s standard protocol. The differentially expressed genes were identified using the DESeq2 tool. Identification of differentially translated genes was performed using the RiboDiff tool.

ChIP-seq libraries were prepared using the ChIP-IT^®^ Express Enzymatic Shearing (Active Motif) and ChIP-seq library preparation (Vazyme) kits. Primary antibodies used for ChIP included RNAPII Ser5P (61,986, Active Motif); RNAPII Ser2P (61,984, Active Motif); PCF11(A303-706A, Bethyl); CPSF160(A301-580A, Bethyl); WDR33(A301-152A, Bethyl); CstF77(A301-094A, Bethyl); CPSF100(A301-583A, Bethyl); CPSF30(A301-585A, Bethyl); Fip1(A301-462A, Bethyl); CFIm68(A301-356A, Bethyl); These libraries were sequenced on the NovaSeq platform, and all ChIP-seq data processing and analysis were performed according to the ENCODE ChIP-seq pipeline (https://www.encodeproject.org/data-standards/chip-seq/). Metagene plots were generated with the deepTools2.0 computeMatrix tool with a bin size of 10 bp. Graphs representing the (IP/Input) signal (ChIP-seq) were then created with R package profileplyr. Metagene profiles are shown as the average of two biological replicates.

iCLIP-seq libraries were prepared following previously reported protocol (16,21,22). The primary antibodies [WDR33 (A301-152A) and CstF64 (A301-092A)] were purchased from Bethyl. Bioinformatics analysis was conducted as previously described (16). Two biological replicates were performed for each iCLIP-seq library, and only the reproducible crosslinking sites were kept for downstream analysis.

## Results

### U1 AMO may affect U1 snRNP structural integrity in vitro

We hypothesized that the 25 nt U1 AMO, routinely used in previous experiments for functional knockdown of U1 snRNA, might disrupt U1 snRNP structural integrity based on two observations. First, in the nuclear extract (NE) prepared from HeLa cells, Co-IP analysis using anti-U1C antibody showed that U1-70K and U1A, two other U1 snRNP specific proteins, were significantly downregulated in the IPed sample using U1 AMO treated NE (23). Second, it is well established that U1-70K can bind U1 snRNA stem loop 1 (SL1) (Figure 1A) (24–26), which is close to the region where 25 nt U1 AMO binds, and U1C was also reported to be associated with U1 snRNA at the 5’ end within U1 snRNP complex (25–27). Therefore, U1 AMO might interfere with the associations of U1 snRNA and U1-70K/U1C.

**Figure 1.**
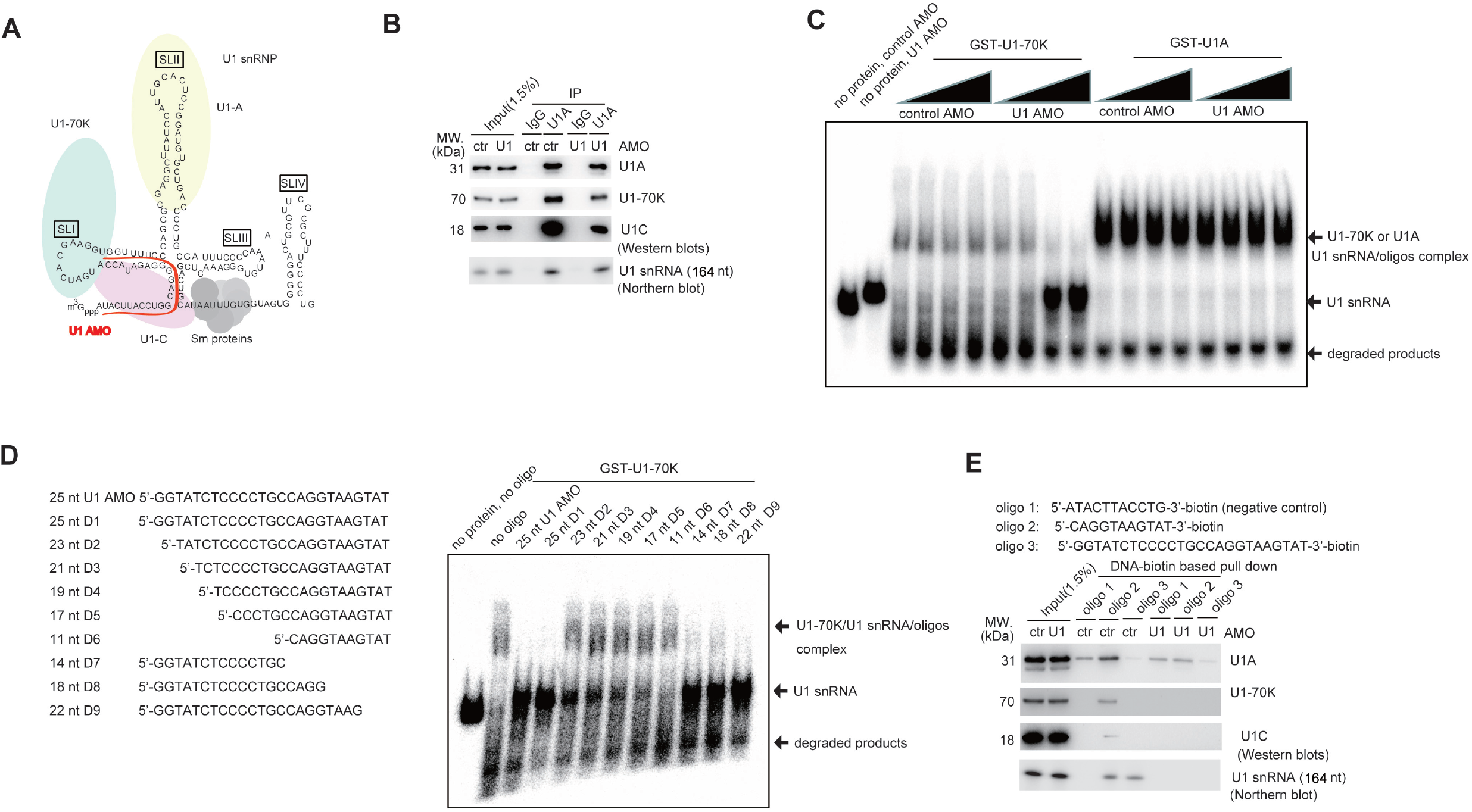
Effect of U1 AMO on U1 snRNP structure in vitro. (A) Schematic representation of components of U1 snRNP complex. The U1 snRNA is illustrated in black and U1 AMO is illustrated in red. U1-specific proteins, including U1-70K (green), U1A (yellow) and U1C(pink), are shown with ellipses. U1-70K and U1A bind SLI and SLII respectively. (B) Co-immunoprecipitation (IP) analysis using antibodies against control IgG or U1A in the presence of control or U1 AMO followed by Western Blotting analysis (for detection of U1-70K, U1A and U1C) and Northern blotting analysis (for detection of U1 snRNA). The input is 1.5% of HeLa NE used for IP. (C) Gel mobility shift assays using detectable amount of radiolabeled U1 snRNA (about 0.2 μM) and recombinant GST-U1-70K (5 μM) or GST-U1A (5 μM) protein in the presence of control or U1 AMO (0, 0.02, 0.2, 2 μM). (D) Gel mobility shift assays using detectable amount of radiolabeled U1 snRNA (about 0.2 μM) and recombinant GST-U1-70K protein in the presence of U1 AMO or DNA oligos. The sequences of the corresponding oligos have been indicated. (E) DNA-biotin based pull down assays using the indicated oligos and HeLa NE followed by Western blotting analysis (for detection of U1-70K, U1A and U1C) and Northern blotting analysis (for detection of U1 snRNA). All experiments were repeated at least three times and representative results are shown.

We first performed a similar Co-IP experiment using HeLa NE to test the above hypothesis. Instead of an anti-U1C antibody, we used an anti-U1A antibody. Consistent with earlier findings, we observed a mild decrease in U1-70K protein and an apparent reduction in U1-C protein in the IPed products for U1 AMO treated samples (Figure 1B). In contrast, similar levels of U1A protein and U1 snRNA were detected. Note that we used a comparable AMO concentration (10 uM) as that before (23), therefore, these experiments could be combined, and the results suggest that U1 AMO might interfere with protein-protein interactions within the U1 snRNP complex in the NE. To test if U1 AMO could affect protein-RNA interactions within the U1 snRNP complex, we took advantage of the knowledge that U1-70K and U1A could directly bind U1 snRNA and performed a gel mobility shift assay in the presence of control and U1 AMO. As expected, recombinant GST-U1-70K and U1A proteins formed stable protein-RNA complexes with U1 snRNA in the presence of control AMO; however, GST-U1C did not form any complex (Supplemental Figure 1A-B). However, we found that U1 AMO could efficiently block U1-70K binding of U1 snRNA even at low concentration (0.2 uM), wherein U1-70K/U1 snRNA was near a 1:1 ratio (Figure 1C). In contrast, the protein-RNA interaction of GST-U1A and U1 snRNA was unaffected in the tested conditions.

As the 25 nt U1 AMO directly binds the 5’ end of U1 snRNA, we predicted that U1 AMO might block U1-70K binding through steric hindrance. We employed a DNA oligo harboring the same sequence with 25 nt U1 AMO, and performed the aforementioned gel shift assay to test this. Indeed, similar to AMO, 25 nt antisense DNA oligo could also efficiently block U1-70K/U1 snRNA association (Figure 1D). The results motivated us to subsequently test the blocking efficiency of a series of truncated forms of U1 antisense DNA. As expected, the shortening of the 5’ end of antisense DNA gradually alleviated the inhibitory effect, which coincided with the fact that the binding region of the 5’ end of antisense DNA was close to U1 snRNA SL1; however, the 3’ end was unaffected (Figure 1D; Supplemental Figure 1C).

To consolidate the above steric hindrance model, we applied a previously reported biotin-based antisense oligo pull-down method to enrich the U1 snRNP complex from HeLa NE (28). The 11 nt biotin-labeled DNA, which is complementary to the free 5’ end of U1 snRNA, could enrich significant amount of U1-70K and U1C in the pull-down assay (Figure 1E). In contrast, the 25 nt biotin-labeled DNA pull-down sample returned undetectable U1-70K and U1C proteins. Northern blotting analysis of U1 snRNA demonstrated that both biotin-DNA oligos could enrich U1 snRNA with similar efficiency (Figure 1E), excluding the possibility that the observed difference resulted from differential U1 snRNA pull-down efficiency. As background controls, we performed the same pull-down assays using NE treated with U1 AMO, and results showed that U1 AMO treatment (10 uM) completely blocked the U1 snRNA 5’ end within NE (Figure 1E). It appears that U1A has some non-specific binding in this assay, while we conclude that at least a portion of U1A was enriched by 11 nt biotin-labeled DNA based on the difference in band intensity.

Based on the above results, we concluded that U1 AMO might significantly affect U1 snRNP complex integrity in the context of HeLa NE.

### U1 AMO may affect U1 snRNP structural integrity in vivo and impact the association between U1 snRNP and RNA polymerase II (RNAPII)

As a first step to explore the molecular mechanisms underlying U1 snRNP telescripting, we performed control and 25 nt U1 AMO transfection in HeLa cells and subsequently prepared NE for the same aformentioned Co-IP analysis using anti-U1A antibody. We and others have shown that transfection of 10 uM U1 AMO is sufficient to induce U1 snRNP telescripting effect and widespread intronic PCPA events at 12 h time point post-transfection in human cells (3,11,29). For the sake of consistency, we used this condition for subsequent U1 AMO transfection experiments. Co-IP and Western blotting analysis revealed that the U1 inhibitory effect is more pronounced than the results obtained in the aforementioned experiment (Figure 1B; Supplemental Figure 2A). This was partly because the nucleus contains less U1-70K and U1C protein upon U1 AMO transfection. To confirm this, we examined their expression in the whole cells, the nucleus, and the cytoplasm, respectively. The results demonstrated that U1 AMO treatment consistently led to an apparent accumulation of U1-70K/U1C/U1A proteins in the cytoplasm and a corresponding downregulation of U1-70K/U1C in the nucleus. In contrast, their overall protein expression was not disturbed (Figure 2A). It must be noted that the observed difference was not caused by potential cross-contamination between cytoplasm and nucleus extracts, as CPSF160, an mRNA 3’ processing factor expressed exclusively in the nucleus, was undetectable in the cytoplasm in both conditions. Additionally, U1 AMO did not appear to cause the U1 snRNA accumulation in the cytoplasm, as revealed by the Northern blotting analysis (Figure 2A).

**Figure 2.**
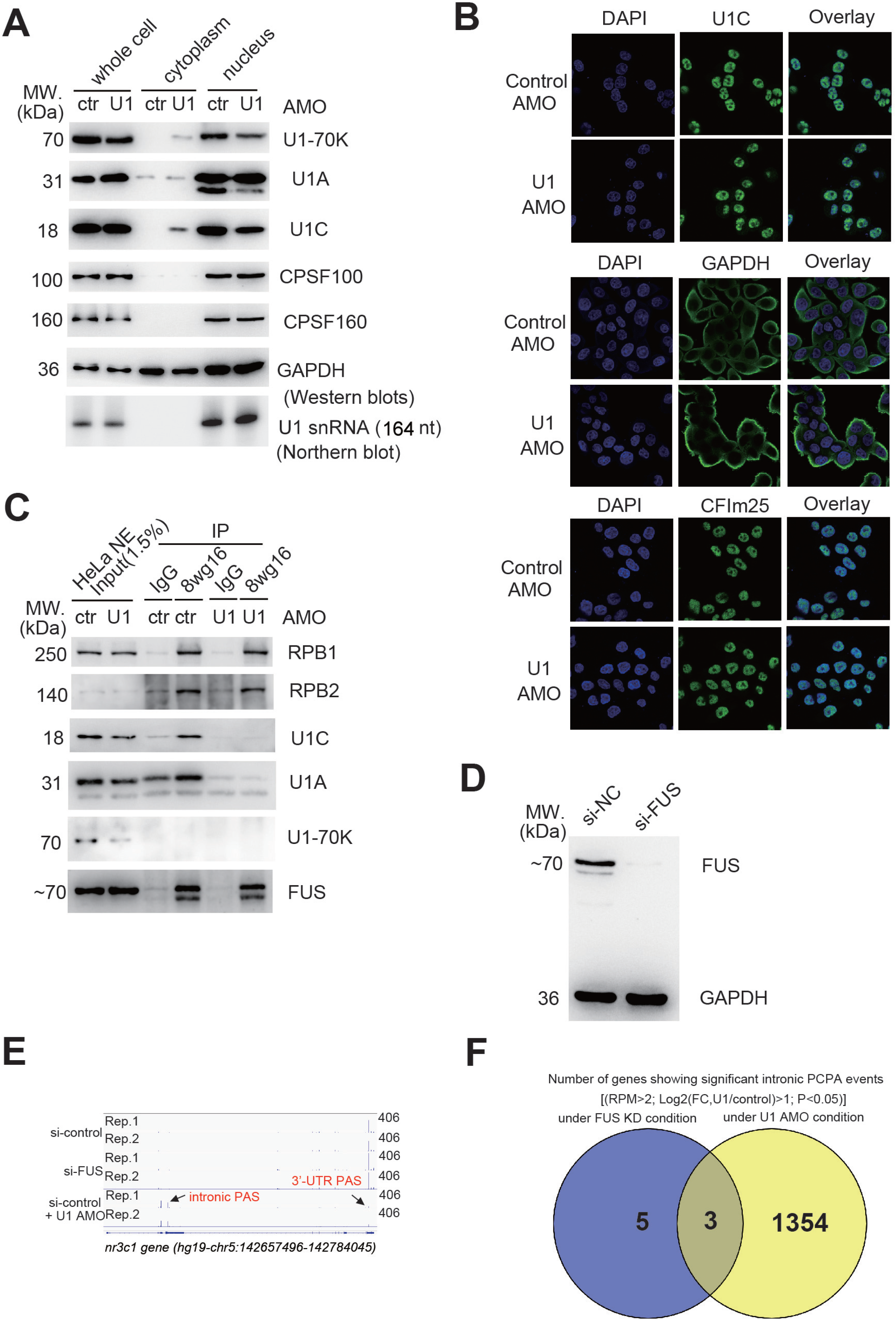
(A) Western blotting analysis (for detection of U1-70K, U1A and U1C) and Northern blotting analysis (for detection of U1 snRNA) to examine the subcellular distribution of U1 snRNP components upon control or U1 AMO transfection in HeLa cells. The indicated proteins serve as controls. (B) Immunostaining of U1C, GAPDH and CFIm25 proteins in control and U1 AMO treated HeLa cells. (C) Co-immunoprecipitation (IP) analysis using control IgG or 8wg16 antibodies in the presence of control or U1 AMO followed by Western blotting analysis (for detection of indicated proteins). The input is 1.5% of HeLa NE used for IP. Experiments were repeated at least three times and representative results are shown. (D) Western blotting analysis of FUS protein in HeLa cells transfected with control or FUS siRNAs. Cells were harvested 72 hour after transfection. (E) IGV track screen shots showing 3’-seq results for *nr3c1* gene in control and FUS siRNA treated HeLa cells. As positive control, U1 AMO was transfected in control cells to detect the intronic PCPA events for *nr3c1* gene. (F) Venn diagram showing the numbers of overlapping and non-overlapping genes that displayed PCPA events using a previously reported pipeline (11).

To complement the results observed in the Western blotting analysis, we performed immunostaining for U1C. A positive immunostaining signal was observed in the cytoplasm for U1 AMO treated cells, while it was exclusively localized in the nucleus for control cells (Figure 2B). These results suggested that U1 AMO transfection significantly affected the overall U1 snRNP structural integrity in vivo. To demonstrate the generality of this phenomenon, we also carried out the above experiments in another human cell line SW480, and the results were the same obtained in HeLa cells (Supplemental Figure 2B-C).

Recent studies have shown that U1 snRNP associates with RNA polymerase II (RNAPII) (23,30–32); we next investigated if this association is disrupted in U1 AMO treated cells. Using a well-established antibody (8wg16) targeting the CTD (C-terminal domain) of RNAPII, our Co-IP analysis confirmed that RNAPII could interact with U1C and/or U1A in the control HeLa cells (Figure 2C). This interaction might be independent of nucleic acids and gene transcription, as shown by the similar results under the treatment of Benzonase or actinomycin D (Supplemental Figure 2D). As expected, this interaction was significantly disrupted in U1 AMO treated HeLa cells (Figure 2C). We failed to detect U1-70K protein in the IPed sample, which might have been lost during the washing steps.

Furthermore, we performed the knockdown (KD) of FUS (Figure 2D). This protein has been reported to mediate the interaction of U1 snRNP between RNAPII and might play a role in U1 telescripting (8,9,23). Subsequently, we performed Co-IP analysis in control and FUS KD NE using 8wg16 antibody. Results suggested that the association of U1 snRNP and RNAPII was unaffected upon FUS KD (Supplemental Figure 2D). Consistently, the 3’-seq analysis revealed that FUS KD did not cause significant PCPA events (Figure 2E-F; Supplemental Figure 2E; Supplemental Table 1), in contrast to the previous study in mouse neuronal cell lines (8). Notably, we used the same siRNA sequence as before (33), and the FUS knockdown efficiency was around 80-90% at both the mRNA and protein levels (Figure 2D; Supplemental Figure 2E). Therefore, we concluded that FUS might not be involved in U1 snRNP telescripting in our experimental condition.

### U1 AMO treatment caused general RNAPII disassociation with chromatin and widespread premature transcription termination (PTT)

Previous RNAPII ChIP-seq analysis from the Dreyfuss laboratory demonstrated that U1 AMO transfection resulted in premature transcription termination (PTT) for all the actively expressed genes in HeLa cells (Supplemental Figure 3A) (3–5), indicating that the interaction of U1 snRNP-RNAPII disrupted by U1 AMO treatment, might have a general impact on gene transcription. To examine this, we utilized an in vitro reconstitution system previously applied to characterize the structure of U1-snRNP/RNAPII (34). In this system, a synthesized pre-mRNA was hybridized to bulged DNA template, mimicking the gene transcription process (Figure 3A). Importantly, U1 snRNP and RNAPII components could be brought together because their binding sites are close to each other within the DNA/RNA hybrid. Instead of using purified U1 snRNP and RNAPII components, we used HeLa NE for in vitro assay. Indeed, by adopting the well-established MS2-tagged RNA affinity purification method (14), U1 snRNP and unmodified RNAPII could be detected in the pull-down sample (Figure 3A). As expected, little U1 snRNP proteins were detected in the presence of U1 AMO. Significantly, decreased pull-down efficiency was observed for RNAPII when U1 AMO was added in the NE compared to the control AMO, as evidenced by Western blotting analysis using antibodies against RPB1 and RPB2 (Figure 3A). These results indicated that RNAPII tended to fall off DNA/RNA hybrid in the presence of U1 AMO. Likewise, RPB1 IP analysis following RNA quantification returned a similar trend (Figure 3A). Taken together, we provided evidence that U1 AMO may directly affect RNAPII association with pre-mRNA in this in vitro assay.

**Figure 3.**
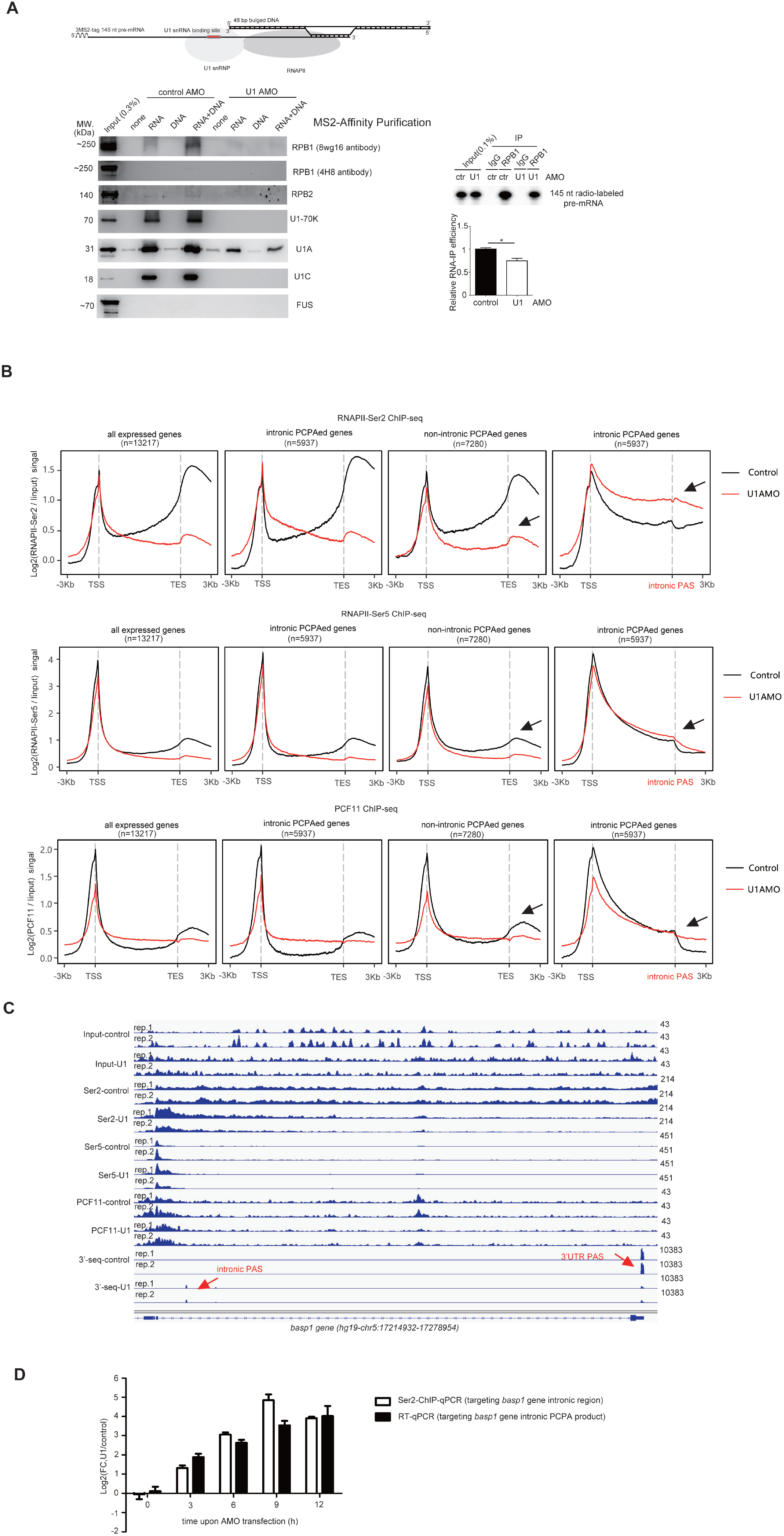
(A) MS2-tagged RNA-based pull-down assay using HeLa NE in the presence of control or U1 AMOs followed by Western blotting analysis to examine the abundance of indicated proteins in the pull-down sample (RNA indicates the 145 nt pre-mRNA; DNA indicates the 48 bp bulged DNA double strand; DNA+RNA indicates that both pre-mRNA and DNA were added in the pull-down assay; none indicates that no nucleic acids were added). The U1 snRNP binding site within the pre-mRNA is labeled red. RNAPII was predicted to bind the DNA/pre-mRNA hybrid. The DNA and pre-mRNA sequences are listed in the Supplemental Table 7. Right picture shows the result of an RNA IP experiment. Radiolabeled pre-mRNA and DNA (cold) were incubated with HeLa NE in the presence of control or U1 AMO. IgG or RPB1 antibodies were used to capture RPB1-associated pre-mRNA (radiolabeled) in the mixture. RNA IP efficiencies were quantified from three independent experiments. (B) Meta-gene plots of RNAPII Ser2P/Ser5P and PCF11 ChIP-seq reads in control and U1 AMO treated HeLa cells for all actively expressed genes (n=13217), intronic PCPAed genes (n=5937), and other non-intronic PCPAed genes (n=7280). For intronic PCPAed genes, a second meta-gene plot was made for each ChIP-seq by replacing the default TES (transcription end sites) with intronic PCPA sites. If multiple intronic PCPA sites were detected for a given gene, the one showing the most significant PCPA was chosen to create the plot. (C) IGV track screen shots showing ChIP-seq(s) and 3’-seq(s) results for *basp1* gene in control and U1 AMO treated HeLa cells. Peaks representing Intronic PAS and canonical PAS are highlighted with red arrow. (D) RNAPII Ser2P ChIP-qPCR and RT-qPCR analysis of *basp1* gene at different time points upon AMOs transfection. The primers are listed in Supplemental Table 7 (ChIP-qPCR primer targets the RNAPII Ser2P binding region upstream of intronic PAS, the input DNA was used as the internal control and for normalization of the ChIP-qPCR data; RT-qPCR primer targets the intronic region to amplify the intronic PCPAed product, *gapdh* gene was used as an internal control and for normalization of the RT-qPCR data). Y-axis represents the log2(fold change) of the results obtained from two samples (U1 versus control AMO). Experiments were performed three times and the standard deviations are shown in error bars.

During in vivo active gene transcription, RNAPII was subjected to different modifications at its CTD repeats, and Ser5P and Ser2P were well-known modifications (35–37). We next asked if U1 AMO treatment in vivo affected the overall Ser5P and Ser2P modifications, along with the unmodified RNAPII. Consistent with earlier findings (38), our Western blotting analysis demonstrated that Ser2P showed a slight decrease in its overall expression upon U1 AMO treatment (Supplemental Figure 3B). This decrease could be significant as it was consolidated by using another antibody phosphorylated forms of RPB1 (Supplemental Figure 3B). We subsequently analyzed their expression in the chromatin fraction, as chromatin is closely associated with gene transcription. The detectable decreases were observed for Ser2P and Ser5P upon U1 AMO transfection, however, unmodified RNAPII was unaffected (Supplemental Figure 3B). As controls, the levels of known 3’ processing factors remained mostly unaltered in corresponding samples. As expected, U1C showed decreased association with chromatin upon U1 AMO treatment. Overall, these results suggested that U1 AMO significantly reduced the level of active RNAPII machinery loaded on chromosomes, which is consistent with the previous result that an overall reduction in RNAPII ChIP-seq signal (unmodified and modified RNAPII) was observed in U1 AMO transfected cells for all actively expressed genes. (Supplemental Figure 3A) (3).

We further performed ChIP-seq analysis for RNAPII Ser5P and Ser2P. Consistent with the previous RNAPII ChIP-seq analysis (3), upon U1 AMO treatment, a progressive decrease in RNAPII Ser5P and Ser2P signals was observed for all the actively expressed genes (Figure 3B; Supplemental Figure 3C; Supplemental Table 2). To gain insight into U1 snRNP telescripting, we divided the actively expressed genes into intronic PCPAed genes and non-intronic PCPAed genes, using our recently published 3’-seq data and QuantifyPoly(A) software (11,20). In total, we found that the usage of 21248 intronic PASs distributed among 5937 genes was significantly elevated upon U1 AMO treatment [Log2 (FC)>1; P<0.05] (Supplemental Table 3). Therefore, intronic PCPAed genes hereafter refer to these 5937 genes, and non-intronic PCPAed genes refer to other 7280 genes (in total 13217 genes), if not indicated otherwise. Indeed, a significant difference between the two subgroups was observed (Figure 3B). For intronic PCPAed genes, a sharp increase in RNAPII Ser2P signal, together with a slight increase in RNAPII and Ser5P signal, were observed between the TSS and intronic PAS, implicating a scenario that RNAPII movement tended to pause toward intronic PAS, thereby providing a time window for intronic PAS processing. For those non-PCPAed genes, RNAPII directly fell off transcribing region in U1 AMO treated cells. Additionally, we performed ChIP-seq analysis for PCF11, a transcription termination-associated factor (39,40), and a similar discrepancy was observed for the two groups of genes (Figure 3B). For intronic PCPAed genes, the PCF11 ChIP-seq signal is stronger upon U1 AMO transfection downstream of intronic PAS, while the opposite trend was observed for non-intronic PCPAed genes. Figure 3C and Supplemental Figure 3D shows each group’s ChIP-seq(s) profiles for several representative genes. Altogether, these data pinpoint a general defect in transcription elongation under the U1 AMO condition. In contrast, a particular RNAPII pausing and high Ser2P modification was observed for intronic PCPAed genes, which was presumed to be associated with mRNA 3’ end processing at the intronic PASs in U1 AMO treated cells.

We further intended to understand whether high RNAPII CTD Ser2P modification occurred before or after PCPA. By focusing on the representative PCPAed gene *basp1*, we found that both PCPA and high Ser2P modifications could occur as early as 3 h time point after U1 AMO transfection, and they are indistinguishable under our experimental condition (Figure 3D). This result aligned with recent reports that Ser2P and mRNA 3’ processing are reciprocally regulated in vivo (41,42).

### Core 3’ processing factors CPSF/CstF play roles in the processing of intronic polyadenylation site (PAS)

As U1 AMO-mediated intronic PCPA is a global effect, we hypothesized that these genes might contain more canonical PASs recognized by cellular 3’ processing machinery during U1 AMO-promoted PTT. To test this, we first confirmed that these intronic PASs share similar canonical PAS patterns as 3’ UTR PASs, including a core A(A/U)UAAA hexamer, a downstream U/GU rich element, and an upstream UGUA motif (Supplemental Figure 4A). By examining the number of core A(A/U)UAAA hexamers, we confirmed that PCPAed genes intrinsically contain more A(A/U)UAAA hexamer motifs in the introns (Supplemental Figure 4B). It must be noted that the values have been normalized to the intronic length, and intronic PCPAed genes are known to have a longer genomic/intronic length than other genes (3). Consequently, we hypothesized that these PCPAed genes are more likely to be recognized by 3’ processing machinery during U1 AMO-triggered PTT.

Furthermore, we wished to identify core 3’ processing machinery associated with the processing of intronic PASs. Although it is predicted that intronic PASs and canonical 3’ UTR PASs share common 3’ processing machinery based on common PAS RNA cis-elements (Supplemental Figure 4A) (11,43), more direct experimental evidence still needs to be provided. To this goal, we adopted a similar RNA affinity approach previously used to purify the 3’ processing complex (14,15). Briefly, three copies of the MS2 hairpin were fused to the intronic PAS of the *nr3c1* gene, which is a bona fide PCPA target upon U1 AMO treatment (Supplemental Figure 3D). To make a negative control RNA substrate, the core hexamer AAUAAA within PAS was mutated to AACAAA (Figure 4A). As expected, this point mutation completely abolished the intronic PAS RNA 3’ processing activity in vitro, as demonstrated by the in vitro cleavage/polyadenylation and in vitro cleavage assays using HeLa NE (Figure 4A). Subsequently, the RNA substrate was incubated with HeLa NE, and the mRNA 3’ processing complex was purified using an MS2 tag. After RNA pull-down, silver staining and quantitative protein analysis by mass spectrometry were performed to estimate the abundance of 3’ processing factors. Indeed, mass spectrometry analysis indicated that most of the top hits were known CPSF/CstF factors (Figure 4B-D; Supplemental Table 4), such as CPSF30, WDR33, CPSF160, Fip1, CstF77, and CstF64, supporting the current model that CPSF/CstF is essential mRNA 3’ processing factors (44–49). Furthermore, we mutated the CstF binding site within the *nr3c1* intronic PAS by converting the ‘U/GU’ rich nucleotides downstream of the core hexamer into ‘A/CA’ nucleotides, and subsequently tested its PAS processing activity in vitro (Sequences were listed in Supplemental Table 7). The results demonstrated that CstF binding played a stimulatory role in intronic PAS processing (Supplemental Figure 4C). To complement the in vitro mRNA 3’ processing assays, we examined the in vivo 3’ processing activity of the above PASs using a luciferase reporter assay that was reported previously (11,16,18,19), and the results were consistent with the in vitro results (Figure 4E). To provide evidence that CPSF/CstF play general roles in intronic PAS processing, we extended the above analysis to the other two PCPAed targets. Indeed, mutations of CPSF/CstF binding sites within the tested PASs decreased the PAS activity by 2 to 10 folds based on the luciferase reporter assay (Supplemental Figure 4D).

**Figure 4.**
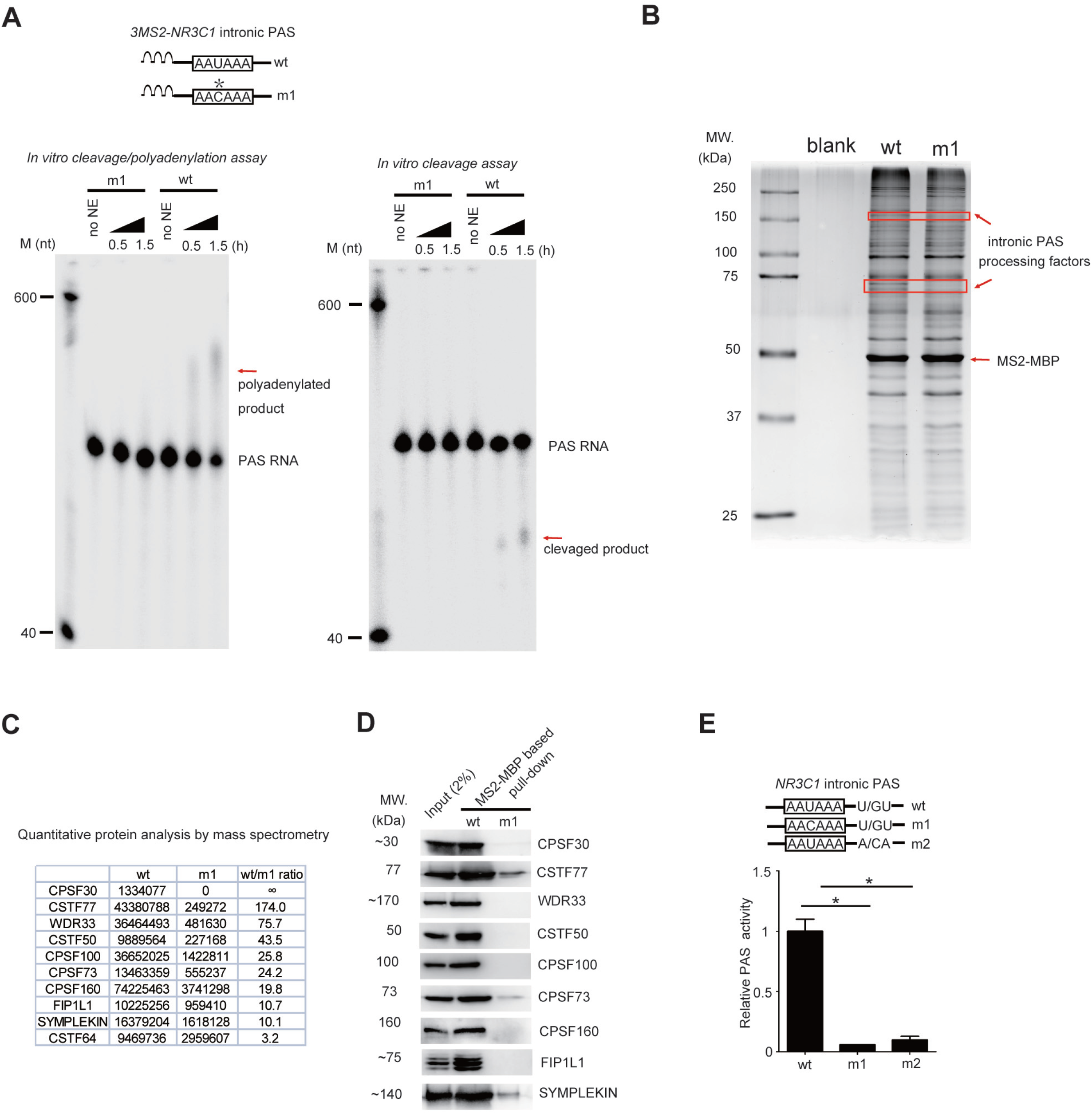
(A) In vitro cleavage/polyadenylation and in vitro cleavage assays using HeLa NE and RNA substrates derived from intronic PAS of *nr3c1* gene. Three repeats of MS2 sequences were added to the 5’ end of the PAS RNA to facilitate the process of protein purification. As negative control, a mutant PAS RNA harboring a point mutation in the core hexamer AAUAAA was prepared simultaneously. The PAS RNA sequences are listed in Supplemental Table 7. The polyadenylated or cleaved RNA products are indicated with red arrows. (B) Silver staining of protein complexes purified from HeLa NE using the indicated PAS RNAs. Potential intronic PAS processing factors, which are indicated with red arrows, could be purified by wt but not m1 PAS RNAs. (C) Results of quantitative protein analysis by mass spectrometry for the indicated proteins in the RNA pull-down sample. (D) Validation of mass spectrometric results by Western blotting analysis using antibodies against indicated proteins. The input is 2% of the total lysates. (E) Measurement of the processing efficiencies of *nr3c1* intronic PAS RNAs and two mutants using pPASPORT system. Experiments were performed three times and the standard deviations are shown in error bars. Student’s t-test was performed to examine the significance of the difference. *P<0.05.

Altogether, our above results provided more direct experimental evidence that known core 3’ processing factors CPSF/CstF were involved in intronic PAS processing upon U1 AMO treatment.

### U1 AMO treatment manipulates CPSF/CstF co-transcriptionally

We further hypothesized that CPSF/CstF associations with transcribed genes might be manipulated under the U1 AMO treatment condition. ChIP-seq(s) were performed to test most of the abovementioned CPSF/CstF factors. Consistent with the current co-transcriptional recruitment model of 3’ processing factors (50,51), a sharp peak at TSS in the ChIP-seq profile for WDR33, CPSF30, CPSF100, FIP1, and CstF77 in control cells was detected. In contrast, another major peak was observed near TES only for WDR33, FIP1, and CstF77 (Figure 5A; Supplemental Figure 5A), coinciding with their roles in canonical PAS processing. Overall, we found that most of the ChIP-seq profiles investigated for CPSF/CstF factors showed significant changes in U1 AMO treated cells (Figure 5A; Supplemental Figure 5A). Among them, CPSF160 showed the most striking change, and it appeared that U1 AMO activated CPSF160 association with genomic regions of both intronic PCPAed and remaining actively transcribed genes. Wdr33, CPSF100, and CstF77 also showed mildly elevated signals across the gene body for all the actively expressed genes. However, for CPSF30 and FIP1, a slight increase in binding signal was only observed for intronic PCPAed genes, whereas the opposite trend was observed for non-PCPAed genes. Altogether, these data suggested that co-transcriptional recruitment of CPSF/CstF was globally enhanced toward intronic PASs upon U1 AMO transfection, which is consistent with the PCPA effects elicited by U1 AMO.

**Figure 5.**
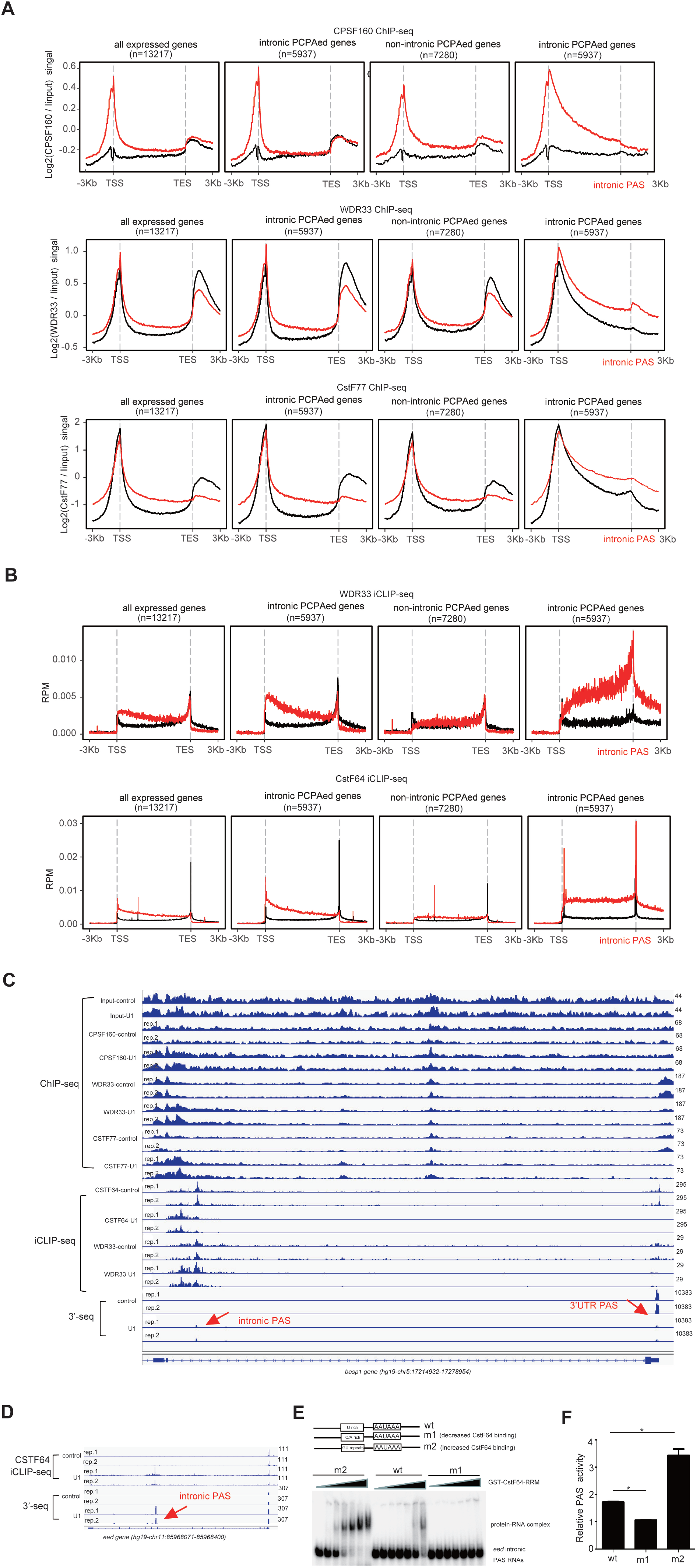
(A) Meta-gene plots of CPSF160, WDR33 and CSTF77 ChIP-seq reads in control and U1 AMO treated HeLa cells for all actively expressed genes (n=13217), intronic PCPAed genes (n=5937), and other non-intronic PCPAed genes (n=7280). For intronic PCPAed genes, a second meta-gene plot was made for each ChIP-seq by replacing the default TES (transcription end sites) with intronic PCPA site. If multiple PCPA sites were detected for a given gene, the one showing the most significant PCPA was chosen to create the plot. (B) Meta-gene plots of WDR33 and CSTF64 iCLIP-seq reads in control and U1 AMO treated HeLa cells for all actively expressed genes (n=13217), intronic PCPAed genes (n=5937), and other non-intronic PCPAed genes (n=7280). For intronic PCPAed genes, a second meta-gene plot was made for each ChIP-seq by replacing the default TES (transcription end sites) with intronic PCPA site. (C) IGV track screen shots showing CPSF160/WDR33/CSTF77 ChIP-seq(s), WDR33/CSTF64 iCLIP-seq(s) and 3’-seq results for *basp1* gene in control and U1 AMO treated HeLa cells. (D) IGV track screen shots showing CSTF64 iCLIP-seq and 3’-seq results for *eed* gene in control and U1 AMO treated HeLa cells. (E) Gel mobility shift assay using recombinant GST-CSTF64-RRM (RNA recognition motif) (0, 1, 2, 5, 10, 15, 20 μM) and indicated PAS RNAs (detectable amount, approximately 0.1 uM). The PAS RNA sequences are listed in Supplemental Table 7. (F) Measurement of the processing efficiencies of *eed* intronic PAS RNAs and two mutants using pPASPORT system. Experiments were performed three times and the standard deviations are shown in error bars. Student’s t-test was performed to examine the significance of the difference. *P<0.05.

To further characterize CPSF/CstF interactions with pre-mRNAs globally before and after U1 AMO treatment, we subsequently mapped protein-RNA interactions in vivo by iCLIP-seq for two canonical CPSF/CstF factors, WDR33 and CstF64, given their direct role in PAS recognition (16,49,52,53). At least three lines of evidence suggested that our iCLIP-seq(s) were reliable. First, meta-gene plots revealed sharp peaks at TES in control cells for both proteins (Supplemental Figure 5B), consistent with their roles in canonical PAS processing. Second, by searching for the most enriched motifs, we found that AAUAAA and U/GU rich motifs were the most significant motifs for WDR33 and CstF64, respectively (Supplemental Figure 5B), consistent with previous reports (16,49). Third, by focusing on the density of the reads closing to the canonical cleavage/polyadenylation (C/P) site, we observed peaks approximately 20 nt upstream of the C/P site for WDR33 and 10 nt downstream of the C/P site for CstF64 (Supplemental Figure 5B), which agree well with the current in vitro model for PAS recognition by CPSF/CstF (16,44,46,54,55). Two major observations were made from the iCLIP-seq analysis between the control and U1 AMO treated cells. First, for intronic PCPAed genes, a significantly elevated iCLIP-seq signal was detected across the gene body for both WDR33 and CstF64 upon U1 AMO treatment (Figure 5B), consistent with the results obtained from ChIP-seq data and suggested the U1 AMO enhanced CPSF/CstF associations with DNA/pre-mRNA co-transcriptionally. Second, for non-PCPAed genes, the change in iCLIP-seq signal value was not as apparent as that in PCPAed genes (Figure 5B), consistent with the data that non-PCPAed transcripts tend to be short and have less PAS motifs recognized by CPSF/CstF (Supplemental Figure 4B). Figure 5C and Supplemental Figure 5C shows several representative examples illustrated by snapshots of mapped reads using the IGV software. We further wished to provide more direct evidence that the enhanced binding of CPSF/CstF across intronic PAS observed in U1 AMO treated cells might directly impact intronic PAS processing (Figure 5B). We were particularly interested in testing the hypothesis that elevated CstF binding signal upstream of intronic PAS might enhance downstream PAS processing based on two observations. Firstly, CstF intrinsically has a broader RNA binding capability than CPSF in vitro (16,44,47–49,53,54,56), and CstF may play a leading role in association with pre-mRNA during co-transcriptional mRNA 3’ processing in vivo. Second, in addition to enhanced CstF64 binding for PCPAed genes downstream of intronic C/P site upon U1 AMO treatment, we also observed a similar pattern upstream of intronic C/P sites (Figure 5B), implying CstF64 binding upstream of C/P sites might also contribute to intronic PAS processing. In this regard, we selected two PCPAed genes harboring significant CstF64 binding signals upstream of intronic C/P sites (Figure 5D; Supplemental Figure 5D). As shown in Figure 5D, CstF64 displayed a higher binding capacity to the target sequence under the U1-AMO treatment. To manipulate differential CstF64 binding stringency at this specific region, we replaced wild-type sequences with strong or weak CstF64 binding motifs, namely ‘GU’ repeats and ‘CA’ rich sequences (Figure 5E; Supplemental Figure 5D; Sequences were listed in Supplemental Table 7), based on previous studies (53,56). Gel mobility shift assays confirmed the relatively strong and weak binding affinities of corresponding sequences with CstF64 (Figure 5E; Supplemental Figure 5D). Indeed, using the aforementioned luciferase reporter assays, we found that strong CstF64 binding upstream of the C/P site returned relatively high PAS activity in both cases. In contrast, the opposite trend was observed for construct harboring low-CstF64 binding motif (Figure 5F; Supplemental Figure 5D). These examples indicated that enhanced CstF binding upstream of intronic PASs induced by U1 AMO might directly contribute to the PCPA effect.

### PCPAed products could be exported to the cytoplasm and translated

We further asked if U1 AMO-induced PCPAed products could be exported to the cytoplasm and translated. In this regard, we extracted RNAs from cytoplasm fraction and total cell extracts and subsequently carried out a 3’-seq analysis. Two nuclear polyadenylated non-coding RNAs, malat1 and neat1, are barely detectable in cytoplasmic RNAs based on the 3’-seq data (Supplemental Figure 6A), suggesting that the nucleus might minimally contaminate the cytoplasmic fractionation. By calculating the export efficiency of the U1-AMO-induced PCPAed products (cytoplasm/total reads ratio), we found that >80% of them could be exported to the cytoplasm using a cutoff value of 0.2 (Figure 6A; Supplemental Table 5). This number is in line with current reports that most of the reported intronic polyadenylated products could be exported to the cytoplasm (29,57).

**Figure 6.**
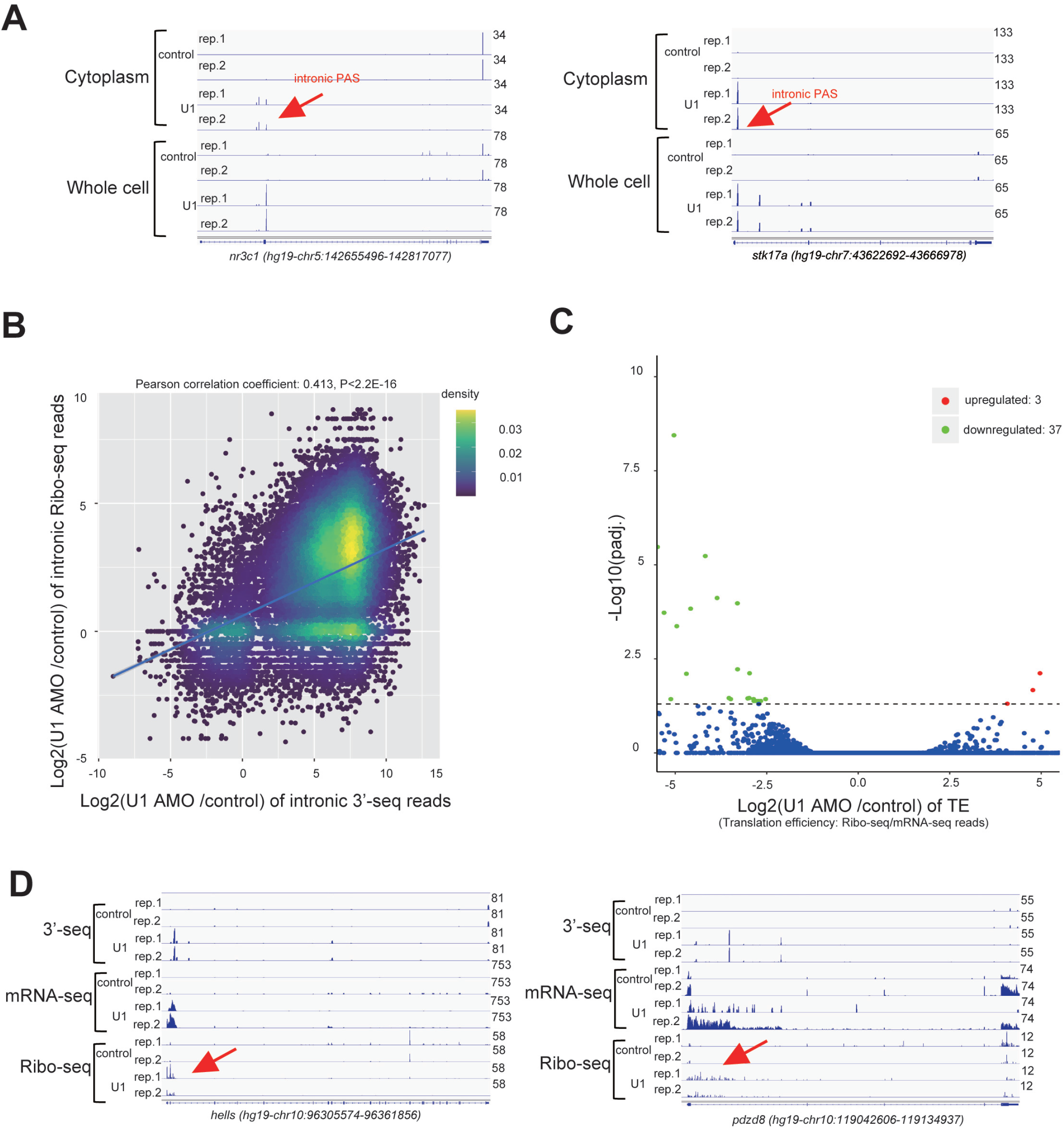
(A) IGV track screen shots showing 3’-seq results for *nr3c1* and *stk17a* genes in cytoplasmic and total RNAs prepared from control and U1 AMO treated HeLa cells. (B) Comparison of the fold changes between intronic Ribo-seq and intronic 3’-seq reads in control and U1 AMO treated samples. (C) Volcano plot showing the differentially translated genes in control and U1 AMO treated HeLa cells (translation efficiency: Ribo-seq reads/mRNA-seq reads). Upregulated genes are indicated red and downregulated genes are indicated green. (D) IGV track screen shots showing Ribo-seq, mRNA-seq and 3’-seq results for *hells* and *pdzd8* gene in control and U1 AMO treated HeLa cells.

To understand if these intronic PCPAed transcripts could be translated, we performed Ribo-seq analysis in control and U1 AMO treated HeLa cells. By assessing the 3-nucleotide (nt) periodicity of Ribo-seq data (Supplemental Figure 6B), we concluded that our Ribo-seq data is of good quality to estimate the translational landscape. Multiple lines of evidence suggested that the majority of these intronic PCPAed products have the potential to be translated. First, consistent with the results from mRNA-seq, Ribo-seq revealed that intronic reads increased by 2 to 9 folds upon U1 AMO transfection (Supplemental Figure 6C-F). Overall, the fold change of intronic PCPAed reads correlates well with intronic Rib-seq reads. (Figure 6B). Second, by calculating the translation efficiency (TE, Ribo-seq/mRNA-seq reads ratio), we found that U1 AMO treatment only affected the TE of 39 genes, among which eight were PCPAed genes (Figure 6C). Third, by comparison of intronic Ribo-seq reads upstream of intronic C/P sites, we found that 58.53% (6744/11523) of introns of the PCPAed genes might be translated upstream of intronic C/P sites [Log2(RPM, FC)>1; P<0.05], which accounted for 71.00% (4215/5937) of the PCPAed genes (Figure 6D; Supplemental Figure 6G). Altogether, consistent with other reports (29,57), we concluded that most of these intronic PCPAed products could be exported to the cytoplasm and have the potential to be translated.

## Discussion

In this study, we provided direct evidence that U1 AMO may significantly disrupt U1 snRNP structure itself, thereby affecting the activity of its associated factor RNAPII, which eventually caused premature transcription termination (PTT) and facilitated the co-transcriptional recruitment of core 3’ processing factors CPSF/CstF to act on cryptic PAS. Additionally, we showed that these PCPAed products might be translated, providing biological significance for U1 snRNP telescripting. A schematic model has been proposed in Figure 7. Overall, our data provided more understanding of the phenomenon of U1 snRNP telescripting and consolidated the currently prevalent model that regulation of transcription and polyadenylation is highly correlated.

**Figure 7.**
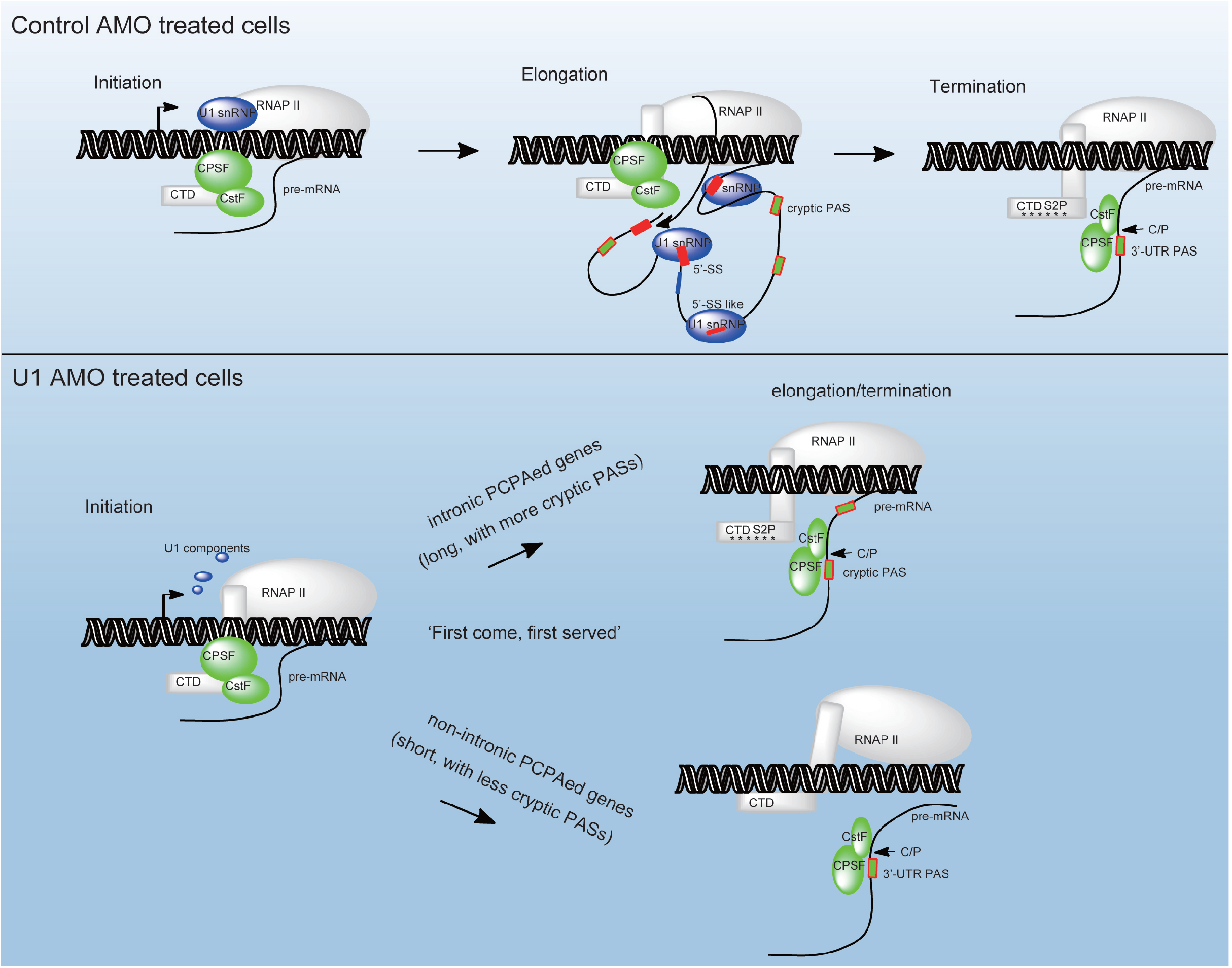
A schematic diagram depicting the model for U1 snRNP telescripting. In addition to splicing, U1 snRNP could be a key player in sustaining transcription elongation by associating with RNAPII in control cells. Although CPSF/CstF could be loaded onto chromatin during the transcription initiation step, they are not fully assembled onto pre-mRNA 3’UTR PAS until RNAPII reaches the end of transcription unit, wherein RNAPII CTD Ser2 is heavily phosphorylated. In U1 AMO treated cells, the integral U1 snRNP complex might fall apart (some of the U1 specific proteins could be exported to the cytoplasm), which affected its association with RNAPII and ultimately affected transcription elongation. Based on the well-accepted ‘first come, first served’ co-transcriptional mRNA 3’ processing model (62,63), the CPSF/CstF complex could be fully assembled onto cryptic PASs of intronic PCPAed genes once U1 AMO-triggered premature transcription termination (PTT) occurs. The intronic mRNA 3’ processing, RNAPII pausing and RNAPII CTD Ser2P could be reciprocally regulated near intronic PAS. In contrast, non-intronic PCPAed genes tend to be short and have less intronic PAS, their 3’ end formation takes place soon after the 3’ UTR PAS is transcribed.

To elucidate the U1 snRNP telescripting mechanism, the previous U1-CPAF activation model has focused on the compositions and functions of U1 snRNP-associated factors (6). Unlike this model, we found that the 25 nt U1 AMO could significantly affect the protein-protein interaction and protein/RNA interaction within the U1 snRNP complex and may directly cause downstream transcription defects and aberrant 3’ processing factors associations with DNA/pre-mRNAs. Notably, our model and the previous U1-CPAF model may not be mutually exclusive. For example, the U1-CPAF complex, which was captured by formaldehyde crosslinking followed by mass spectrometry analysis, is dynamic and transient in nature. In our experimental condition, U1 AMO treatment only moderately affected the abundance of U1 snRNP-specific proteins in the nucleus (Figure 2A). In addition, not all the molecular interactions within U1 snRNP were affected, such as the interaction of U1A and U1 snRNA (Figure 1C). Therefore, U1-CPAF might exist in the form of U1-specific protein/CPAF in the nucleus. Consistent with the U1-CPAF model, our data implied that the concentration of CPAF, or so-called core 3’ processing factors, near intronic PAS, were significantly elevated in the U1-AMO treatment condition (Figure 5B).

Our study has also raised several interesting questions for future investigation. First, RNAPII ChIP-seq data from the Dreyfuss laboratory and our RNAPII Ser2P ChIP-seq analysis have demonstrated that transcription dynamics are also globally changed for non-PCPAed genes upon U1 AMO treatment (Figure 3B) (3). In contrast, RNA-seq analysis revealed that the overall level for most of those non-PCPAed genes was not affected (Supplemental Table 6). It will be interesting to understand the discrepancy between RNAPII ChIP-seq and mRNA-seq data, as it provides insight into understanding the potential reciprocal regulations between transcription and pre-mRNA metabolism. Secondly, consistent with current co-transcriptional mRNA 3’ end processing model (41,42,50), RNAPII pausing and high RNAPII Ser2P density were observed near the intronic PAS for those intronic PCPAed genes upon U1 AMO treatment, while the trend was not apparent for non-PCPAed genes (Figure 3B), indicating that Ser2P and 3’ end processing might be uncoupled during the process of transcription termination for these non-PCPAed genes under U1 AMO condition, which adds a layer of complexity to the regulatory mechanism of co-transcriptional mRNA 3’ end processing. Third, although our study and previous studies have suggested that the U1 snRNP and RNAPII might have physical interactions (Figure 2C) (23,30–32), it remains unclear if this association is direct or indirect, particularly in a cellular context, which may enable us to understand how U1 AMO caused the differential RNAPII activity. Fourth, although we have provided data that U1 AMO disrupts U1 snRNP structure and affects transcription elongation, which emerges an important factor in regulation of alternative polyadenylation (APA) (58–59), we still do not know how U1 protects against intronic PAS usage, and whether it associates with U1’s role in splicing needs to be further investigated, given the intimate links between 3’ processing, splicing and transcription regulation (37,57,60,61). Finally, it will be interesting to characterize the protein outputs of the PCPAed transcripts. Although we provided evidence that these PCPAed products could be translated in cells, more direct evidence is still required.

## Accession Numbers

All the deep sequencing data have been deposited to GEO database with the accession no. GSE192943.

## Acknowledgments

We thank Dr. Marc-Étienne Huot for providing pGEX-6P2-U1A, pGEX-6P2-U1C and pGEX-6P2-U1-70K plasmids.

## Figure Legends for Supplemental Figures

Figure S1. (A) Commassie blue staining of purified recombinant GST-U1-70K, GST-U1A, GST-U1C and GST-CstF64-RRM (RNA recognition motif) proteins. Red arrow in each lane indicates the band of the expressed target protein. (B) Gel mobility shift assays using detectable amount of radiolabeled U1 snRNA (about 0.2 μM) and GST protein or recombinant GST-U1-70K (5 μM), GST-U1C (5 μM) protein or GST-U1A (5 μM) in the presence of control or U1 AMO (2 μM). (C-D) Replicates of Figure 1D.

Figure S2. (A) Co-immunoprecipitation (IP) analysis were performed using NE prepared from HeLa cells transfected with control or U1 AMO, and antibodies against control IgG or U1A. Western blotting analysis (for detection of U1-70K, U1A and U1C) and Northern blotting analysis (for detection of U1 snRNA) were subsequently carried out to determine the relative Co-IP efficiency. The input is 1.5% of HeLa NE used for IP. (B) Western blotting analysis (for detection of U1-70K, U1A and U1C) and Northern blotting analysis (for detection of U1 snRNA) to examine the subcellular distribution of U1 snRNP components upon control or U1 AMO transfection in SW480 cells. The indicated proteins serve as controls. (C) Immunostaining of U1C, GAPDH and CFIm25 proteins in control and U1 AMO treated SW480 cells. (D) Co-immunoprecipitation (IP) analysis using control IgG and 8wg16 antibodies under various conditions (ActD treatment; Benzonase treatment; si-U1A/U1C/U1-70K/FUS treated) followed by Western blotting analysis (for detection of indicated proteins). The input is 2% of HeLa NE used for IP. (E) IGV track screen shots showing 3’-seq results for *fus* gene, *gapdh* gene, and *cmip* gene in control, FUS siRNA, and U1 AMO treated HeLa cells.

Figure S3. (A) Meta-gene plots of RNAPII and normalized Ser5P/Ser2P (normalized to RNAPII) ChIP-seq reads in control and U1 AMO treated HeLa cells for all actively expressed genes (n=13217), intronic PCPAed genes (n=5937), and other non-intronic PCPAed genes (n=7280). For intronic PCPAed genes, a second meta-gene plot was made for each ChIP-seq by replacing the default TES (transcription end sites) with intronic PCPA site. RNAPII ChIP-seq data were downloaded from Dreyfuss publication (3). (B) Western blotting analysis of indicated proteins in corresponding subcellular fractions (whole cell extracts, chromatin, nucleus and cytoplasm) prepared from HeLa cells treated with control or U1 AMO. (C) MA-plots showing the significantly differential binding sites in the presence of U1 AMO based on the original ChIP-seq data. Peak calling was performed using MACS3 software and DiffBind package was used to identify the differential binding events. The number of decreased affinity sites and increased affinity sites are shown in the plot. (D) IGV track screen shots showing ChIP-seq and 3’-seq results for PCPAed genes (*cmip, nr3c1*) and non-PCPAed genes (*gapdh, rps11*) in control and U1 AMO treated HeLa cells.

Figure S4. (A) Density plot showing the frequencies of ‘A(A/U)UAAA’, ‘U/GU’ and ‘UGUA’ elements near the indicated PAS groups. (B) Boxplots showing the distribution of gene size/intronic size/number of A(A/U)UAAA/normalized number of A(A/U)UAAA of intronic PCPAed genes (n=5937) and non-intronic PCPAed genes (n=7280). Boxplot midlines represent the median, box limits span the first and third quartiles. Two-samples Wilcoxon rank sum test was used to examine the significance of the difference. *P<2.2e-16. (C) In vitro cleavage/polyadenylation and in vitro cleavage assays using HeLa NE and RNA substrates derived from intronic PAS of *nr3c1* gene. To create CstF64 binding mutant, downstream U/GU rich sequences were replaced with A/CA sequences. The PAS RNA sequences are listed in Supplemental Table 7. (D) Measurement of the processing efficiencies of b*asp1* and *cmip* intronic PAS RNAs and two mutants using pPASPORT system. Experiments were performed three times and the standard deviations are shown in error bars. Student’s t-test was performed to examine the significance of the difference. *P<0.05.

Figure S5. (A) Meta-gene plots of CPSF100, CPSF30, FIP1L1, and CFIm68 ChIP-seq reads in control and U1 AMO treated HeLa cells for all actively expressed genes (n=13217), intronic PCPAed genes (n=5937), and other non-intronic PCPAed genes (n=7280). For intronic PCPAed genes, a second meta-gene plot was made for each ChIP-seq by replacing the default TES (transcription end sites) with intronic PCPA sites. If multiple PCPA sites were detected for a given gene, the one showing the most significant PCPA was chosen to create the plot. (B) Sequence logo based on all reproducible crosslinking nucleotides and 50 nt (WDR33)/10 nt (CSTF64) on each side. MEME (Multiple EM for Motif Elicitation) of the top 1000 WDR33/CSTF64 binding sites. The most enriched motifs from each group are shown. (C) IGV track screen shots showing CPSF160/WDR33/CSTF77 ChIP-seq(s), WDR33/CSTF64 iCLIP-seq(s) and 3’-seq results for PCPAed genes (*cmip* and *nr3c1*) and non-PCPAed genes (*gapdh* and *rps11*) in control and U1 AMO treated HeLa cells. (D) Gel mobility shift assay using recombinant GST-CSTF64-RRM (RNA recognition motif) (0, 1, 2, 5, 10, 15, 20 μM) and indicated PAS RNAs (detectable amount, approximately 0.1 uM). The underlined sequences were used for gel shift assay. The bar graphs are the measurement of the processing efficiencies of *rrm2* intronic PAS RNAs and two mutants using pPASPORT system. Experiments were performed three times and the standard deviations are shown in error bars. Student’s t-test was performed to examine the significance of the difference. *P<0.05.

Figure S6. (A) IGV track screen shots showing 3’-seq results for *malat1*, *neat1* and *gapdh* genes in cytoplasmic and total RNAs prepared from control and U1 AMO treated HeLa cells. (B) Distribution of 5’-end of 30 nt Ribo-seq reads near start codon and stop codon regions. 2 biological replicates were performed for each sample (control and U1 AMO treated). (C) Distributions of genomic regions of mapped clean mRNA-seq from control and U1 AMO treated HeLa cells (2 biological replicates). (D) Statistics of Ribo-seq reads (2 biological replicates). (E) Sequence length distribution of clean Ribo-seq reads (2 biological replicates). (F) Distributions of genomic regions of mapped clean Ribo-seq data from control and U1 AMO treated HeLa cells (2 biological replicates). (G) Statistics of ribosome protected fragment (RPF) in introns with intronic PAS. RPM (reads per million) represents normalized RPF.

