## Supplementary figures and images for "U1 AMO (antisense morpholino oligo) disrupts U1 snRNP structure to promote intronic premature cleavage and polyadenylation (PCPA)"

### Supplemental Figure 1

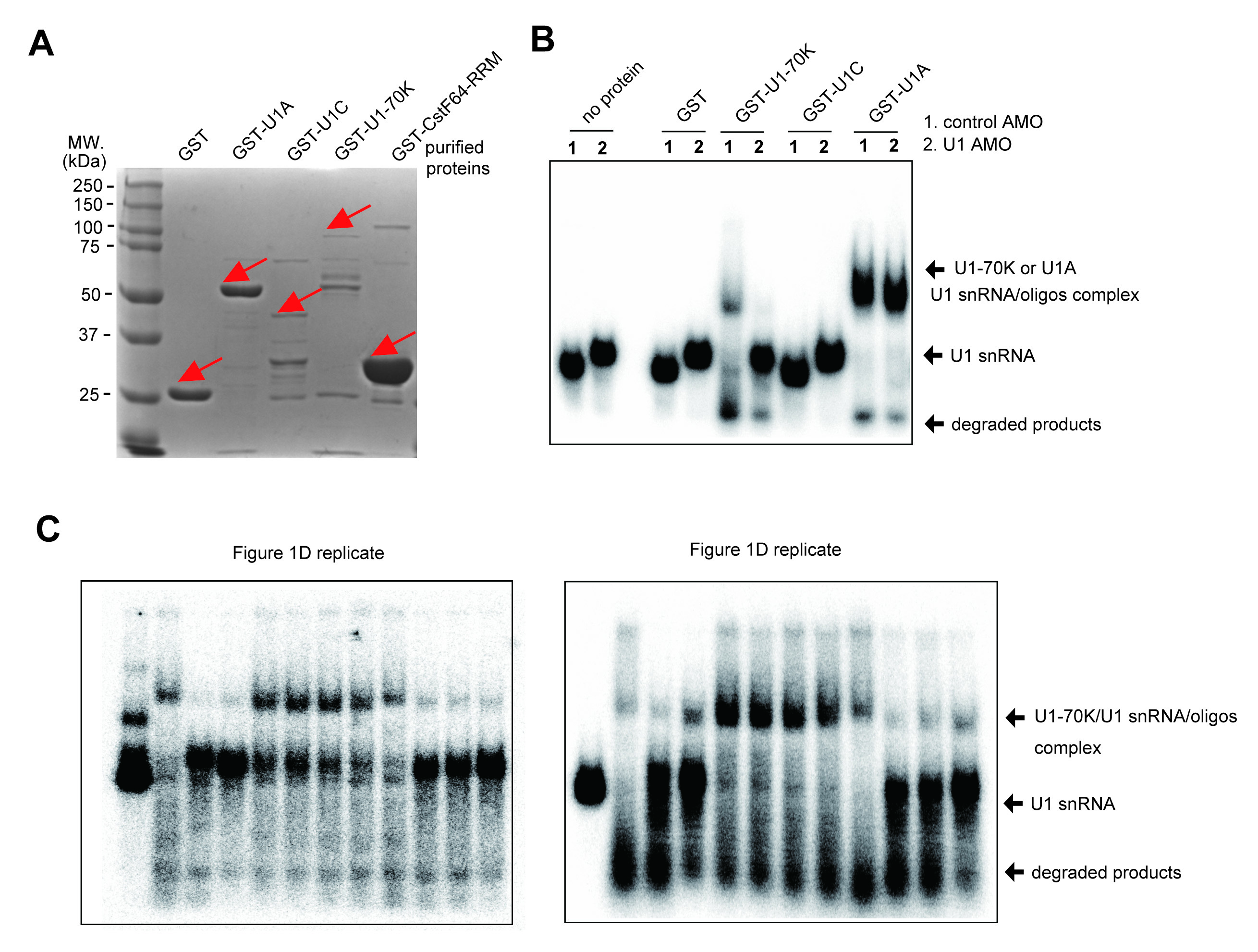

### Supplemental Figure 2

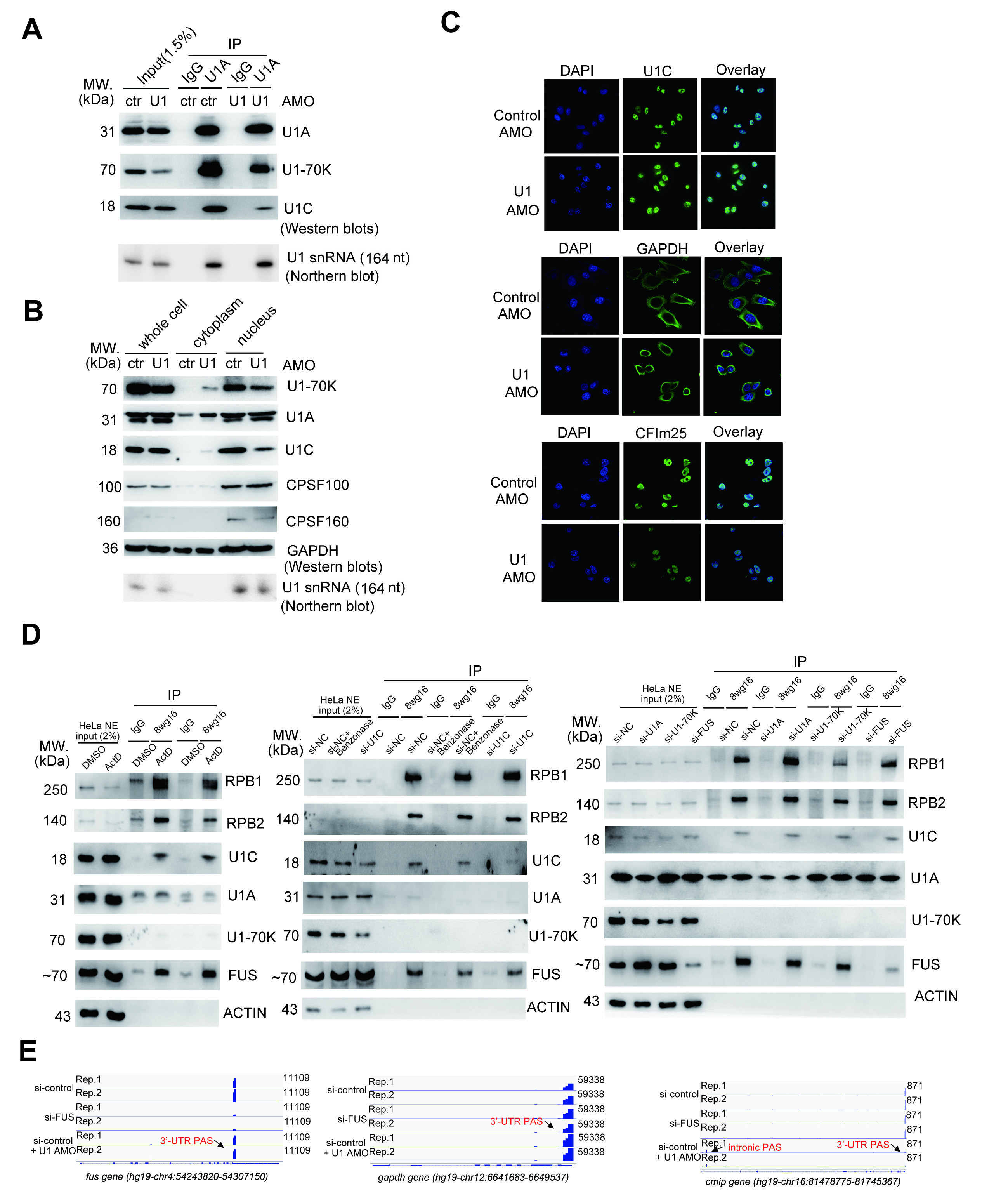

### Supplemental Figure 3

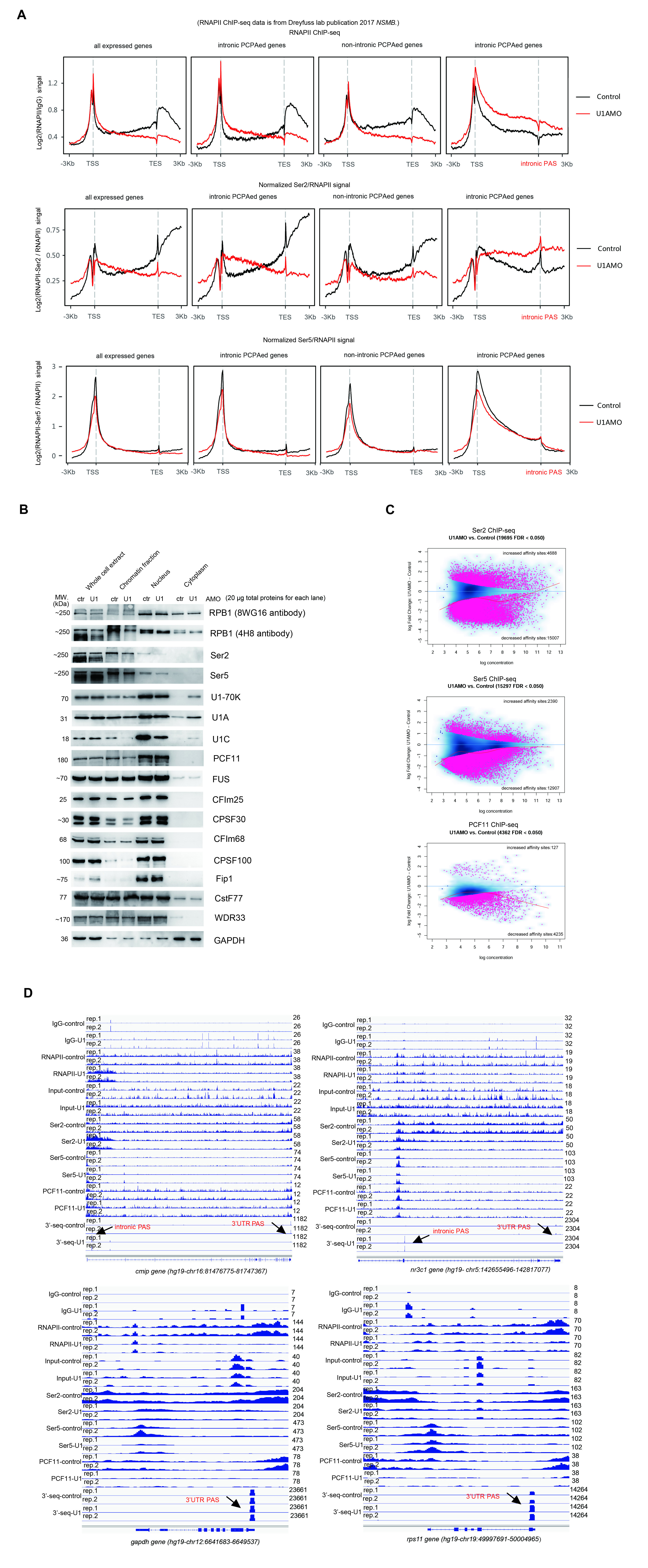

### Supplemental Figure 4

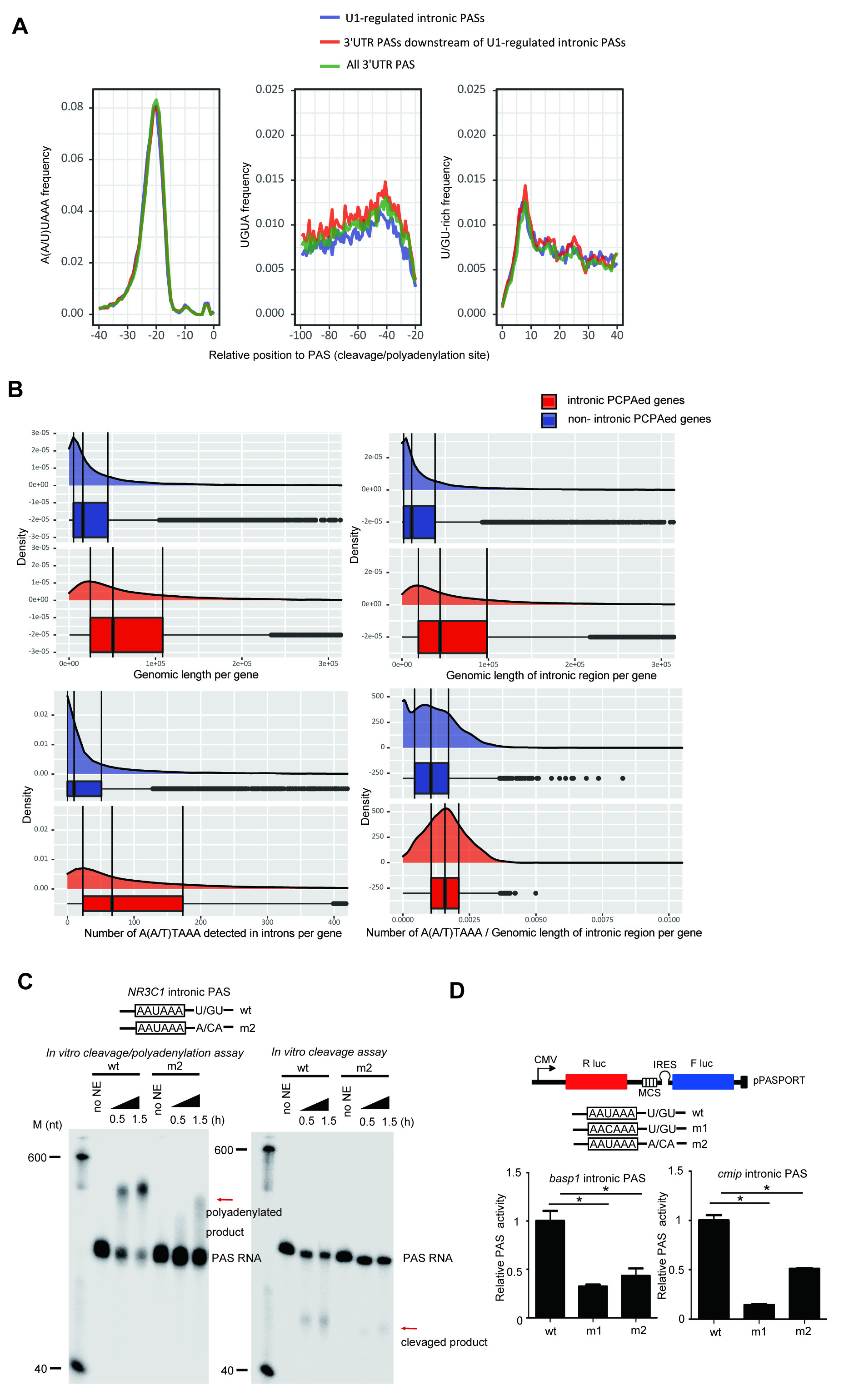

### Supplemental Figure 5

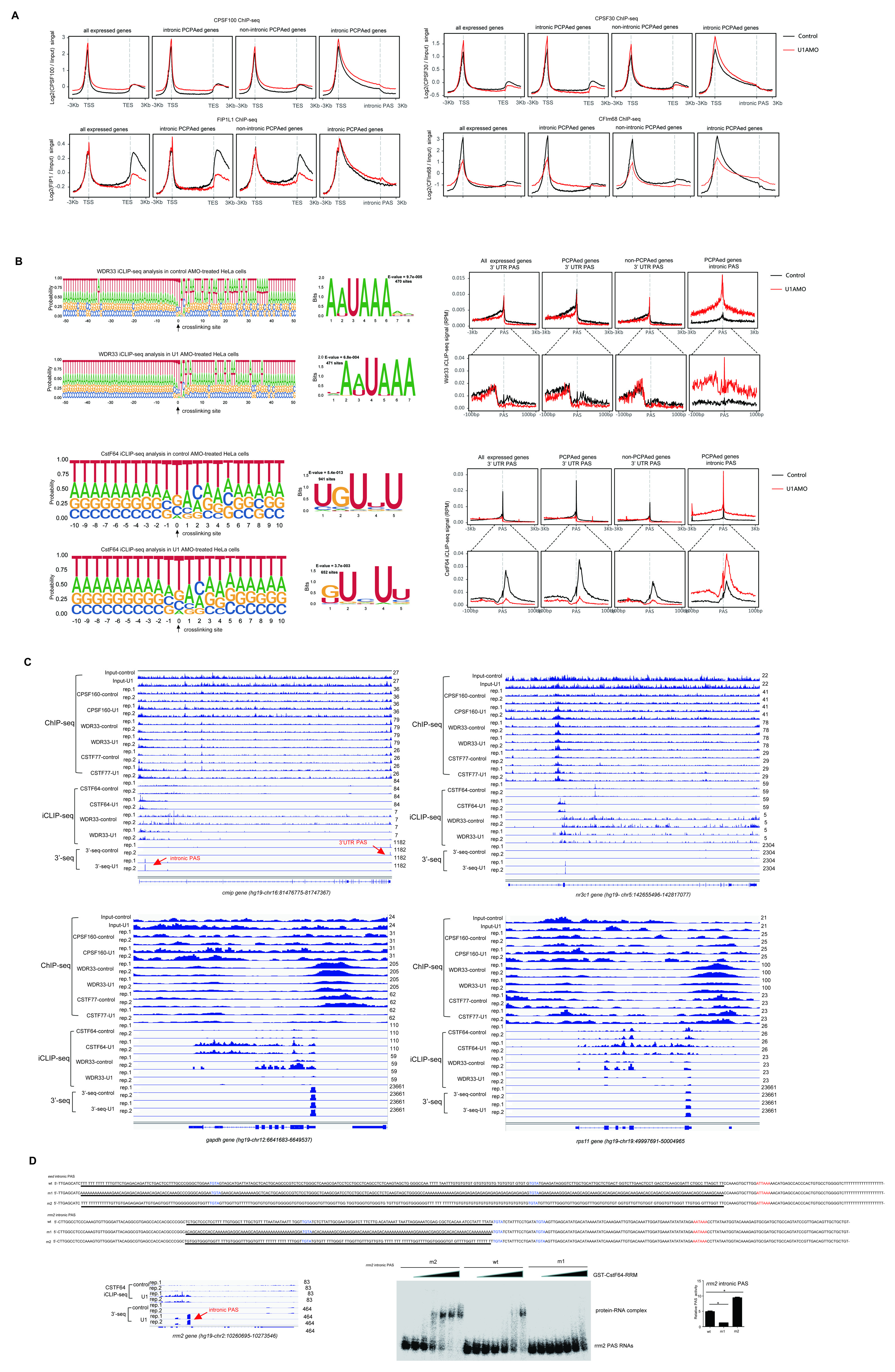

### Supplemental Figure 6

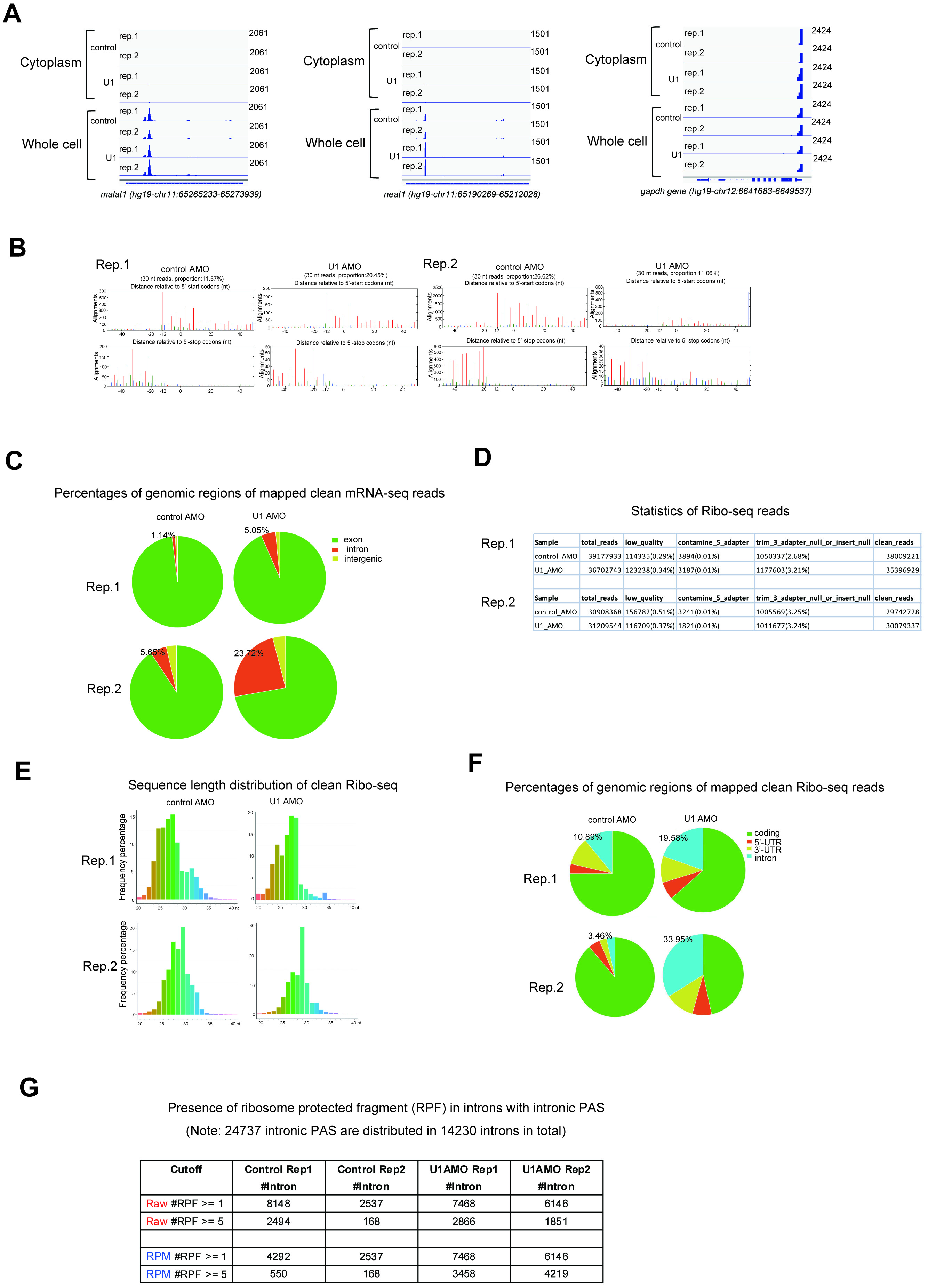
